# Rapid and reversible dissolution of biomolecular condensates using light-controlled recruitment of a solubility tag

**DOI:** 10.1101/2024.01.16.575860

**Authors:** Ellen H. Brumbaugh-Reed, Kazuhiro Aoki, Jared E. Toettcher

## Abstract

Biomolecular condensates are broadly implicated in both normal cellular regulation and disease. Consequently, several chemical biology and optogenetic approaches have been developed to induce phase separation of a protein of interest. However, few tools are available to perform the converse function—dissolving a condensate of interest on demand. Such a tool would aid in testing whether the condensate plays specific functional roles, a major question in cell biology and drug development. Here we report an optogenetic approach to selectively dissolve a condensate of interest in a reversible and spatially controlled manner. We show that light-gated recruitment of maltose-binding protein (MBP), a commonly used solubilizing domain in protein purification, results in rapid and controlled dissolution of condensates formed from proteins of interest. Our optogenetic MBP-based dissolution strategy (OptoMBP) is rapid, reversible, and can be spatially controlled with subcellular precision. We also provide a proof-of-principle application of OptoMBP, showing that disrupting condensation of the oncogenic fusion protein FUS-CHOP results in reversion of FUS-CHOP driven transcriptional changes. We envision that the OptoMBP system could be broadly useful for disrupting constitutive protein condensates to probe their biological functions.

## Introduction

Many proteins are organized into biomolecular condensates in either normal or diseased cell states^1–4^. Biomolecular condensates form as a result of many weak, multivalent interactions, leading to co-assembly of proteins and/or nucleic acids into a dense phase containing a high concentration of the condensed components surrounded by a dilute phase^5^. Often, protein phase separation is facilitated by proteins’ intrinsically disordered regions (IDRs): amino acid sequences which lack a fixed secondary structure^6^ and associate with one another through electrostatic, cation-pi, pi-stacking, and hydrophobic interactions^7^. Biomolecular condensates are thought to play important roles in a diverse array of biological processes such as stress responses^8^, immune regulation^9^, genome organization^10^, and transcription^11,12^. Condensates are thought to carry out biological functions by acting as organizational hubs for biochemical processes, either by colocalizing interacting partners to increase reaction rates or by sequestering components to inhibit their ability to interact^13^. Furthermore, aberrant phase transitions are indicated in a number of pathologies ranging from cancer to neurodegenerative disorders^13–15^.

Due to the importance of phase separation in the regulation of biological processes, a wide variety of optogenetic and chemogenetic strategies have been developed to trigger phase separation of engineered proteins. For example, the optoDroplet system consists of an IDR fused to the blue-light sensitive Cry2^PHR^ protein domain^16^. Upon light stimulation, Cry2 oligomerizes and nucleates condensates which grow through IDR-driven interactions. Several additional tools rely on a similar molecular logic, either through constitutive fusion of an IDR to a light-switchable oligomerization domain (e.g., PixELLs)^17^ or through light-dependent association between an IDR and a constitutive oligomerization domain (e.g., Corelets and CasDrop)^18,19^. Alternative architectures have also been explored, such as light-induced binding between proteins containing multiple discrete folded interaction domains (e.g., the iPOLYMER system)^20^ or the inducible removal of a solubility tag from a phase-separating IDR (e.g., the SPLIT system)^21,22^.

Yet while there are numerous techniques to induce phase separation, comparatively few tools exist to reverse this process by driving naturally phase-separated proteins from a clustered to a dispersed state. One historically important technique to dissolve biomolecular condensates relies on the addition of the organic compound 1,6-hexanediol to inhibit weak hydrophobic interactions^10,23–25^. However, the compound’s indiscriminate activity means it cannot be used to specifically dissolve a condensate of interest, and recent studies suggest that it may also interfere with many other cellular processes (e.g., enzyme activity)^26,27^. Insertion of a protease cut-site can be used to remove IDRs from a protein upon protease addition, resulting in dissolution of the protein clusters^21^. However, cleavage and removal of IDR sequences may disrupt other IDR-encoded biological functions (e.g., short linear motifs for binding, localization, or post-translational modification)^28^. The DISCO system is a promising recent approach^29^ in which rapamycin-inducible dimerization between FRB and FKBP^30^ is used to drive dissolution of a condensate-forming protein, yet the mechanism and efficacy of this approach are still incompletely understood. Moreover, none of these approaches are compatible with rapid or spatially precise control, as protease cleavage is irreversible and dissociation of rapamycin-induced dimers only occurs over hours after rapamycin washout^31^.

Thus, there is still an unmet need for tools to dissolve biomolecular condensates in the complex environment of the cell. Ideally, such a tool should rapidly disperse a specific endogenous condensate of interest without cleaving or otherwise altering the sequences of its constituents. The tool should also be reversible and spatially controlled, enabling dissolution of condensates at desired times and subcellular locations. This level of precision could be used to study particular condensates within a cell, such as those specifically associated with a single transcriptionally-active gene locus^32^. Finally, such a strategy should ideally be broadly applicable to target any endogenous condensate-forming protein of interest.

Here, we report progress toward these goals by developing an optogenetic strategy for IDR-based condensate dissolution. Our approach centers on light-based recruitment of maltose binding protein (MBP), a widely-used purification tag that has been used to increase solubility of a broad range of proteins during bacterial expression and purification, including proteins which undergo phase separation, *in vitro* and *in vivo*^21,22,33–35^. We show that in addition to its utility as a fusion tag, MBP can be transiently recruited to a target protein to dramatically and reversibly alter its partitioning between soluble and condensed phases. Condensates can be manipulated with precise temporal and spatial control, enabling targeted dissolution of individual condensates within the nucleus. We further show that light-based MBP recruitment acts on condensates across several proteins including all three FET family members (FUS, EWS, and TAF15), an RGG domain, and the RNA-binding protein Oskar, suggesting that this approach may be of broad utility for interrogating consequences of protein phase separation. Finally, we apply our OptoMBP approach to probe an exemplary functional context: transcription induced by condensates of the oncogenic transcription factor FUS-CHOP. We show that light-based dissolution is sufficient to partially revert aberrant FUS-CHOP gene expression within 6 hours, suggesting a direct role of FUS-CHOP phase separation in gene expression. Our approach opens the door to rapid, reversible, and precise control over protein organization in both native biological processes and disease states.

## Results

### Recruitment of a solubility tag to biomolecular condensates results in dissolution

The propensity of a protein to undergo phase separation can be regulated by many factors including concentration^36–38^, temperature^39,40^, salt concentration^21^, pH^41^, post-translational modifications^42,43^, and interactions with additional proteins^44,45^. We reasoned that temperature, salt concentration and pH are nonspecific, post-translational modifications can have many consequences beyond tuning phase separation, and that it is difficult to alter total protein concentration rapidly and with high subcellular precision. We thus focused on tuning phase separation through binding to candidate “solubilizing” protein domains.

As a first step toward this goal, we established a model condensate that could be expressed and screened for binding-induced dissolution. Our model condensate was based on the intrinsically disordered region (IDR) of Fused in Sarcoma (FUS)^46–48^, a sequence that has been studied extensively as part of prior optogenetic and chemogenetic tools for manipulating biomolecular condensates^16–18^. At typical cellular concentrations, a single copy of the FUS^IDR^ remains dispersed unless it is linked to additional oligomerization domains^16^. We hypothesized that a protein containing multiple FUS^IDR^ sequences would condense without the need for additional oligomerization domains, forming a simple IDR-driven constitutive condensate in cells. Indeed, a construct containing three FUS^IDR^ repeats, the FusionRed fluorescent protein, an iLID optogenetic recruitment domain^49^, and an FRB chemogenetic recruitment domain^30^ (3xFUS^IDR^-FusionRed-FRB-iLID; **Fig. 1A**) constitutively formed protein droplets in the nucleus when transfected into HEK293T cells (**Fig. 1B**). This construct is compatible with both optogenetic and chemogenetic dimerization systems through the interactions between iLID-SspB and FRB-FKBP, respectively^49,30^.

**Fig. 1.**
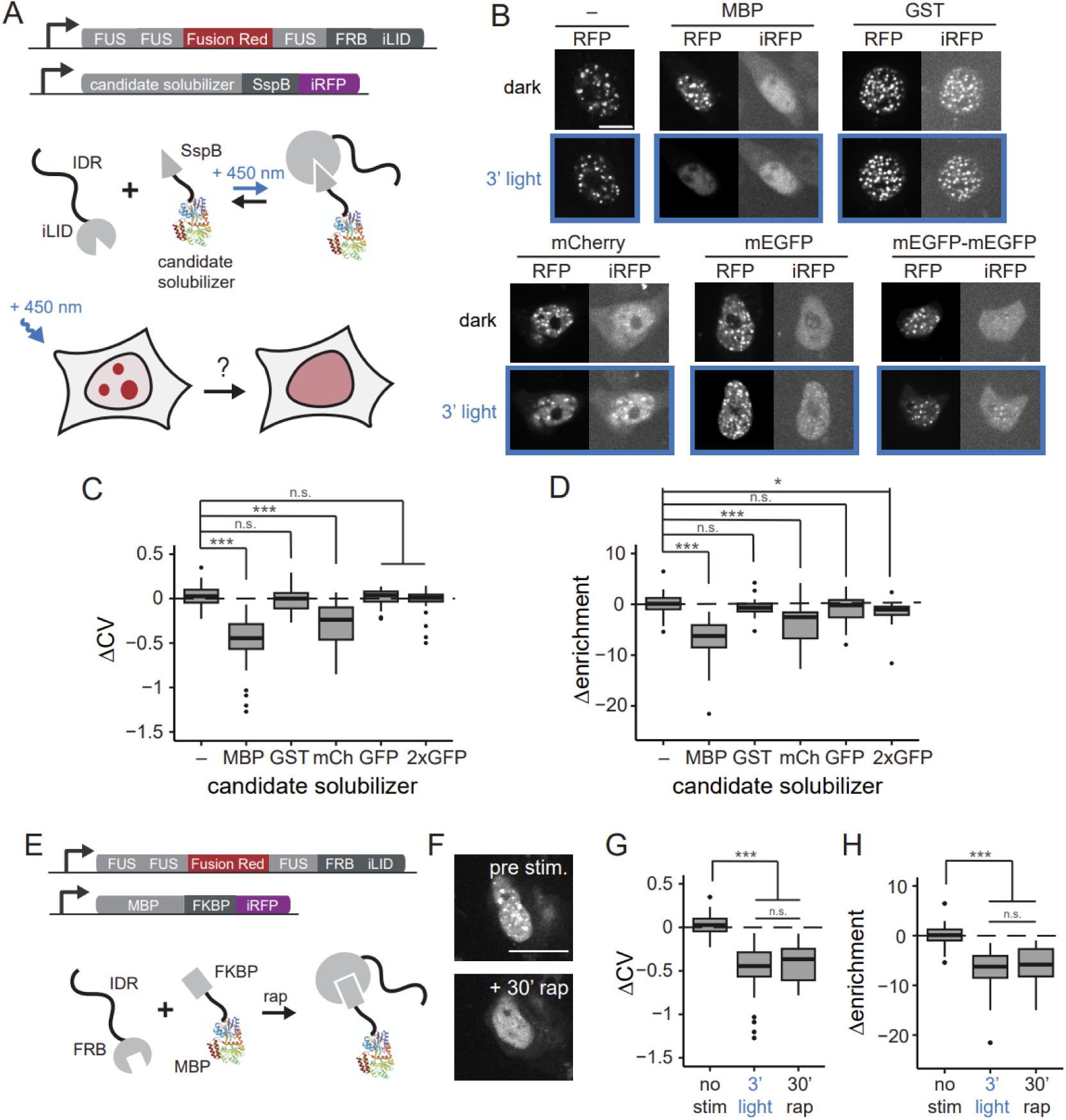
Screen for light controlled dissolution of FUS^IDR^ based condensates. **A.** Schematic of approach. A fusion protein consisting of three copies of the FUS IDR forms stable droplets when expressed in HEK293T cells. We brought candidate solubilizing proteins to 3xFUS^IDR^ droplets using the blue-light sensitive dimerization pair, iLID/SspB. **B.** Representative images showing changes in 3xFUS^IDR^ (RFP channel) and candidate dissolver protein (iRFP), expressed in HEK293T cells, before and after 3 min stimulation with blue light. **C.,D.** Quantification of dissolution efficiency from microscopy data by calculating the change in the coefficient of variation (std/mean) and enrichment (max/mean) of RFP pixels (3xFUS^IDR^) in the nucleus after blue light stimulation. Negative values indicate condensate dissolution. **E.** Schematic of approach for chemogenetic dissolution. FRB/FKBP dimerize in response to rapamycin. **F.** Representative images showing changes in 3xFUS^IDR^ after 30 min rapamycin treatment. **G.,H.** Quantification of dissolution efficiency from microscopy data by calculating the change in the coefficient of variation and enrichment. Dissolution efficacy from blue optogenetic MBP-dimerization was compared to chemogenetic MBP-dimerization. Scale bars: 10 μm.

We next set out to determine whether optogenetic recruitment of a solubilizer domain could drive dissolution of 3xFUS^IDR^ condensates. We chose MBP as a lead candidate, a protein commonly used to increase the solubility of proteins purified from bacterial expression systems^50^. The mechanism of MBP-induced solubilization is still an active area of research^51,52^, but occurs both in cells and *in vitro*^22^, suggesting that a direct interaction between MBP and a target protein is sufficient to mediate solubilization. MBP has also been shown to prevent condensate formation during protein purification of a wide variety of IDRs^33–35^. We also tested Glutathione S-transferase (GST), an affinity tag commonly used in protein purification that has also been suggested to promote solubility^53^, and mCherry, which was previously suggested to inhibit protein phase separation^29^. Finally, we also tested single and double repeats of mEGFP to probe if condensation dissolution might be a function of cargo protein size. Each candidate domain was fused to SspB, which forms one-to-one heterodimers with iLID upon blue light illumination, along with iRFP for visualization. We used non-fluorescent mutants of mCherry^54^ and mEGFP^55^ in our candidate optogenetic dissolution systems so as not to interfere with the detection of other fluorescent proteins in the assay.

We co-transfected HEK293T cells with our 3xFUS^IDR^ construct and a candidate SspB- and iRFP-tagged solubilizer domain and imaged cells before and after treatment with 3 min of 450 nm blue light to dimerize SspB and iLID (**Fig. 1B**). Qualitatively, we observed no illumination-induced changes in 3xFUS^IDR^ condensates in cells expressing no solubilizer construct, GST, mEGFP, or mEGFP-mEGFP. GST colocalized to FUS^IDR^ condensates even prior to blue light exposure, indicative of a constitutive, light-independent interaction between these constructs that did not inhibit FUS^IDR^ condensate formation. We observed weak light-induced dissolution upon mCherry recruitment. Notably, only MBP recruitment appeared to drive complete dissociation of 3xFUS^IDR^ droplets upon illumination.

To quantify the results of these experiments, we selected cells for analysis with comparable expression levels of FUS^IDR^ and the solubilizer protein (**Fig. S1A-B**) and quantified the changes in the coefficient of variation (CV) and local enrichment (enrichment) of FusionRed pixel intensity in the cell nucleus upon light illumination (**Fig. 1C-D**). Cells with a high degree of FUS^IDR^ condensate formation have a high CV in pixel intensity, as they harbor both bright pixels within condensates and dark pixels between them. We estimated the FusionRed enrichment in condensates by dividing the maximum FusionRed pixel intensity by the mean pixel intensity in the nucleus. Both CV and enrichment are expected to decrease following condensate dissolution. We only observed a significant change in both metrics for MBP and mCherry recruitment, with a larger magnitude of effect for MBP (**Fig. 1C-D**), results that are consistent with our qualitative observations (**Fig. 1B**). Notably, the extent of dissolution was not correlated with the size of the recruited solubilizer domain as measured by its computed radius of gyration^56^ (**Fig. S1C**). Because each domain was recruited using the same optogenetic binding motif, we can conclude that the extent of dissolution is primarily determined by properties other than cargo size, concentration, or binding affinity.

We performed additional analyses to further characterize MBP-driven condensate dissolution. We benchmarked our results to a gold standard by comparing the extent of MBP-induced dissolution to 1,6-hexanediol-induced dissolution, a chemical that has been frequently used to disperse a variety of condensates^10,23–25^. We observed similar changes in FusionRed CV and enrichment upon both hexanediol treatment and light-induced MBP recruitment, confirming our qualitative observation that both treatments appeared to drive complete dissolution of the 3xFUS^IDR^ condensates (**Fig. S1D-F**). We also implemented a chemogenetic variant of the MBP recruitment system that takes advantage of the FRB domain on our 3xFUS^IDR^ construct for rapamycin-based protein dimerization^30^ (**Fig. 1E**). We transfected the 3xFUS^IDR^ construct with either the SspB-tagged or FKBP-tagged MBP construct and selected cells with similar expression levels for analysis (**Fig. S1G-H**), which revealed comparable dissolution of 3xFUS^IDR^ condensates in response to either 3’ of illumination or 30’ of rapamycin treatment (**Fig. 1F-H**). Together, these results suggest that inducible MBP recruitment acts as a potent dissolver of FUS^IDR^ condensates. Based on its superior performance in these early screens, we focused on light-based MBP recruitment for subsequent experiments, which we termed the OptoMBP system.

### Optogenetic condensate dissolution is rapid, reversible, and spatially controlled

We next set out to map the spatial and temporal precision of OptoMBP-induced condensate dissolution. To characterize the dynamics of disassembly, we imaged 3xFUS^IDR^ and MBP localization in living cells at high time resolution after blue light illumination (**Fig. 2A**). We quantified the localization of both constructs over time using the coefficient of variation, i.e. the standard deviation of pixel intensity divided by mean pixel intensity in each cell’s nucleus, as reported previously^17^. Quantification across multiple cells revealed that MBP was recruited into FUS droplets within ∼5 sec after blue light illumination (**Fig. 2B**), matching previously-measured kinetics of optogenetic recruitment in living cells^57,58^. The resulting 3xFUS^IDR^/MBP condensates subsequently dispersed with a characteristic timescale of ∼13 sec, comparable to the kinetics observed in the PixELL system, where light is used to disrupt interactions between two engineered proteins (FUS^IDR^-PixD and FUS^IDR^-PixE) to drive their disassembly^17^ (**Fig. 2C**). These results indicate that MBP-induced condensate dissolution is exceptionally rapid, a result that is consistent with the highly dynamic nature of FUS^IDR^ condensates^16^ and which is suggestive of a direct effect of MBP association on condensate stability.

**Fig. 2.**
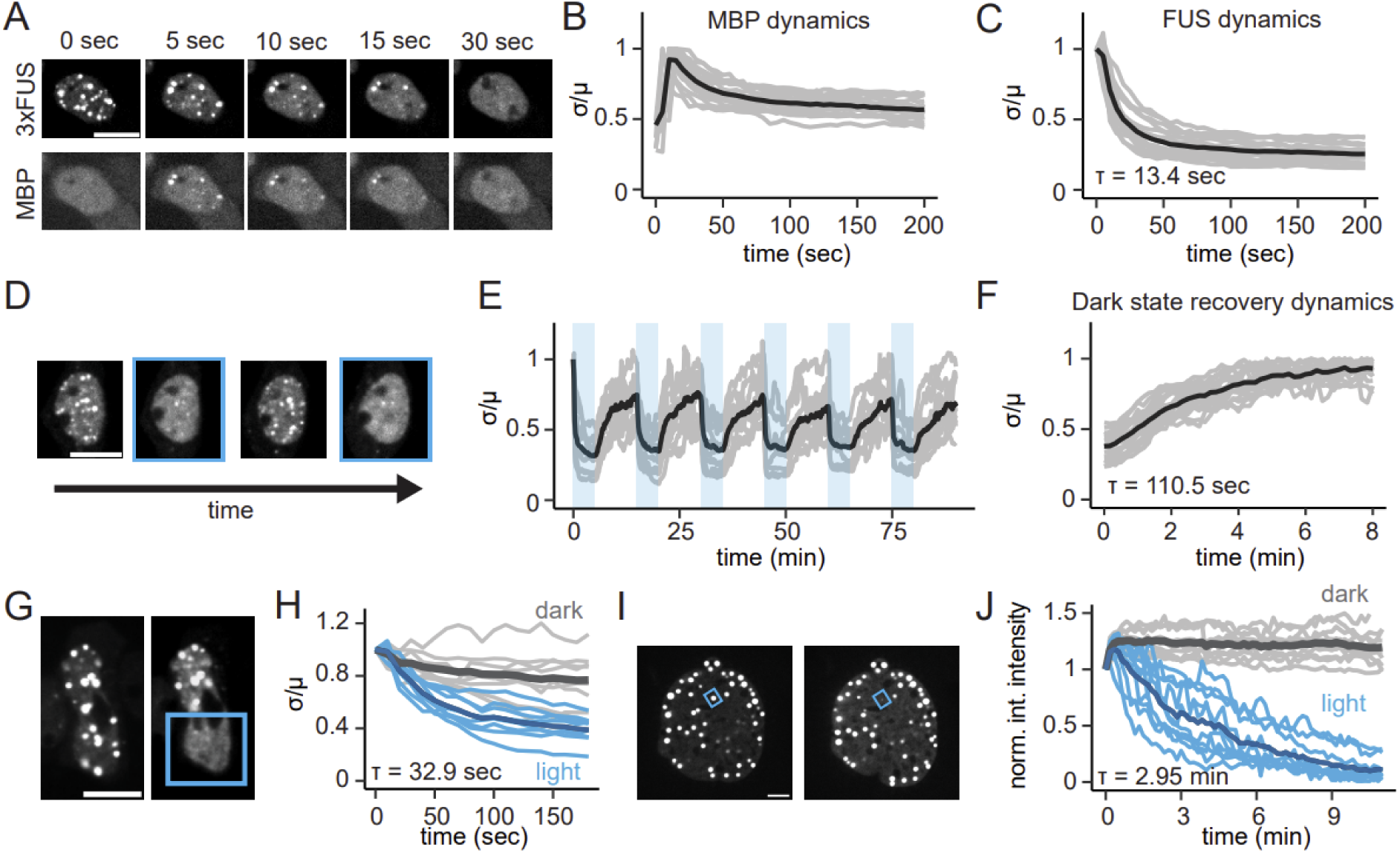
Characterization of temporal and spatial control of condensate dissolution. **A.** Representative time course images showing dissolution 3xFUS^IDR^ condensates and MBP after the addition of blue light (time 0) to dimerize 3xFUS^IDR^ with MBP in HEK293T cell nuclei. MBP rapidly colocalizes with 3xFUS^IDR^ condensates after the addition of blue light. **B.,C.** Dynamics of condensate dissolution were measured by calculating the normalized CV over time. Lines show individual cells with the average shown in bold. For MBP, the CV initially increases indicating colocalization to condensates, then decreases concurrent with condensate dissolution. For 3xFUS^IDR^, the CV decreases rapidly indicating condensate dissolution. Data were fit to the generalized logistic function to calculate a decay rate. **D.** Representative images showing 3xFUS^IDR^ condensate dissolution and formation when cycling between lit and dark states. **E.** Quantification of droplet dissolution reversibility by calculating the normalized CV over time. Blue bars indicate blue light stimulation. **F.** Dynamics of dark state recovery of condensates was measured by calculating the normalized CV over time after removing blue stimulation. **G.** Representative images displaying condensate behavior after blue light illumination is applied to one half of a cell nucleus. Droplets preferentially dissolve in the illuminated region of the cell indicated by the blue box. **H.** Quantification of subcellular dissolution by calculating the CV in the illuminated and non-illuminated regions of the cell.**I.** Images showing illumination and subsequent dissolution of a single FUS-based droplet in a stable NIH3T3 cell line. **J.** Single droplet dissolution was quantified by measuring the integrated pixel intensity of the illuminated droplet over time. Illuminated droplets were compared to non-illuminated droplets several microns away from the illuminated region. Initial increases in pixel intensity were expected due to the increase in FusionRed brightness upon blue light stimulation. Scale bars: 10 μm.

We next tested whether MBP-induced dissolution of 3xFUS^IDR^ condensates was also reversible after removal of the light stimulus, enabling multiple cycles of condensate assembly and disassembly. We switched HEK293T cells expressing 3xFUS^IDR^ and the OptoMBP system between illuminated and dark conditions and measured condensation over time. This experiment revealed that cycles of illumination could drive repeated dissolution and reassembly of 3xFUS^IDR^ droplets with similar kinetics in each cycle (**Fig. 2D-E, Video S1**). Droplet formation in the dark was slower than droplet dissolution in the light, occurring with a characteristic timescale of ∼110 seconds (**Fig. 2F**). This is consistent with prior observations of minutes-timescale FUS^IDR^ condensation in cells^17^ and is likely driven by the timescale of condensate nucleation. Overall, our kinetic data demonstrate that OptoMBP based dissolution of FUS^IDR^-based condensates is both rapid and reversible in living cells.

An additional benefit of optogenetic control is the ability to apply inputs with high spatial precision to nucleate or dissolve droplets at particular positions within the cell^16,17^. To test whether the OptoMBP system is capable of subcellular spatial control, we first illuminated half of an OptoMBP-expressing cell and monitored the spatial distribution of 3xFUS^IDR^ condensates. We observed localized dissolution that was predominantly confined to the illuminated area, even after minutes of continuous illumination (**Fig. 2G-H, Video S2**). We also tested whether the spatial asymmetry persisted after removal of the light stimulus as predicted by prior work^17^. Indeed, condensates in the previously illuminated region remained smaller than droplets in the non-illuminated area several minutes after recovery, confirming that local disassembly produces long-term spatial asymmetries in condensate location (**Fig. S2**).

To further probe the limits of spatial control, we tested whether it might be possible to dissolve a single FUS droplet within the nucleus, a scale that is beyond the reach of prior optogenetic dissolution approaches^17^. Such a finely-targeted stimulus could be useful for perturbing an individual transcriptional locus or other localized condensate-associated process. We conducted this experiment in NIH3T3 cells stably expressing an optogenetically targetable FUS-CHOP oncoprotein (see **Fig. 5** and detailed description below), because the larger nucleus and distance between droplets facilitated illumination of only a single droplet. Indeed, we observed robust dissolution of illuminated single droplets with only minimal effects on droplets in distal regions of the nucleus (**Fig. 2I, Video S3**). Quantification of localized dissolution in both 3xFUS^IDR^ expressing HEK293T cells (**Fig. 2H**) and FUS-CHOP expressing NIH3T3 cells (**Fig. 2J**) revealed a notably slower dissolution timescale of 30 sec - 3 min compared to our previous global illumination experiments (**Fig. 2C**), which may be a consequence of slower photoconversion at the lower illumination intensities that are required for locally precise patterning (see **Methods**) or of diffusion of dark-state 3xFUS^IDR^ molecules from the non-illuminated region maintaining condensates for longer time periods.

### Dissolution efficiency is dependent on solubilizer concentration

How does the efficacy of the OptoMBP system depend on the expression levels of MBP and the target protein (in this case, the 3xFUS^IDR^)? Answering this question may shed light on the mechanism of MBP-induced disassembly. The answer is also of practical importance because it is critical to know whether light-induced condensate disassembly is sensitive to the concentration of system components. In one conceptual model, MBP would act stoichiometrically, so that proportionally more MBP would be required to dissolve increasing levels of FUS. However, intracellular biomolecular condensates are complex and may involve interactions between many components: co-condensing proteins and RNAs^38^, chaperones that regulate condensate assembly and disassembly^59^, or cooperative interactions between multiple monomers (e.g. to nucleate a new condensate), and we reasoned that OptoMBP-based dissolution may also exhibit a complex dose response if MBP acts cooperatively or recruits additional effectors. We thus set out to quantify light-induced condensate dissolution across a range of FUS and MBP expression levels.

We engineered a NIH3T3 mouse fibroblast cell line in which both the 3xFUS^IDR^-FusionRed-FRB-iLID and MBP-SspB-iRFP constructs were stably expressed using lentiviral vectors (**Fig. 1A**). Lentiviral transduction can drive a wide range of protein expression levels depending on integration site, and we sorted cells to cover a ∼10-fold range of expression of both components of the system (**Fig. S3A**). We then imaged single cells to measure their initial FusionRed and iRFP expression as well as the extent of FUS^IDR^ condensation before and after blue light illumination. For dark-incubated cells we observed a sharp threshold in cellular FUS^IDR^ expression above which nuclear condensates were observed and corresponded to high enrichment scores (maximum divided by mean nuclear pixel intensity) (**Fig. 3A**, purple dots). However, while the presence or absence of condensates was well predicted by 3xFUS^IDR^ concentration, we observed that the dilute phase 3xFUS^IDR^ concentration did not plateau at a well defined saturation concentration, as would be expected for a simple single-component condensate, but instead continued increasing with total nuclear 3xFUS^IDR^ levels, a phenomenon that has been previously described for multi-component biomolecular condensates^38^ (**Fig. S3B**). We also observed variable enrichment scores between cells even at the same overall cellular 3xFUS^IDR^ concentration, suggesting that the 3xFUS^IDR^ concentration within phases was variable between cells even at the same overall 3xFUS^IDR^ expression level (**Fig. 3A**). Overall, these data are consistent with 3xFUS^IDR^ condensation existing as a complex process that likely involves interaction with additional components whose levels may vary between cells^60^.

**Fig. 3.**
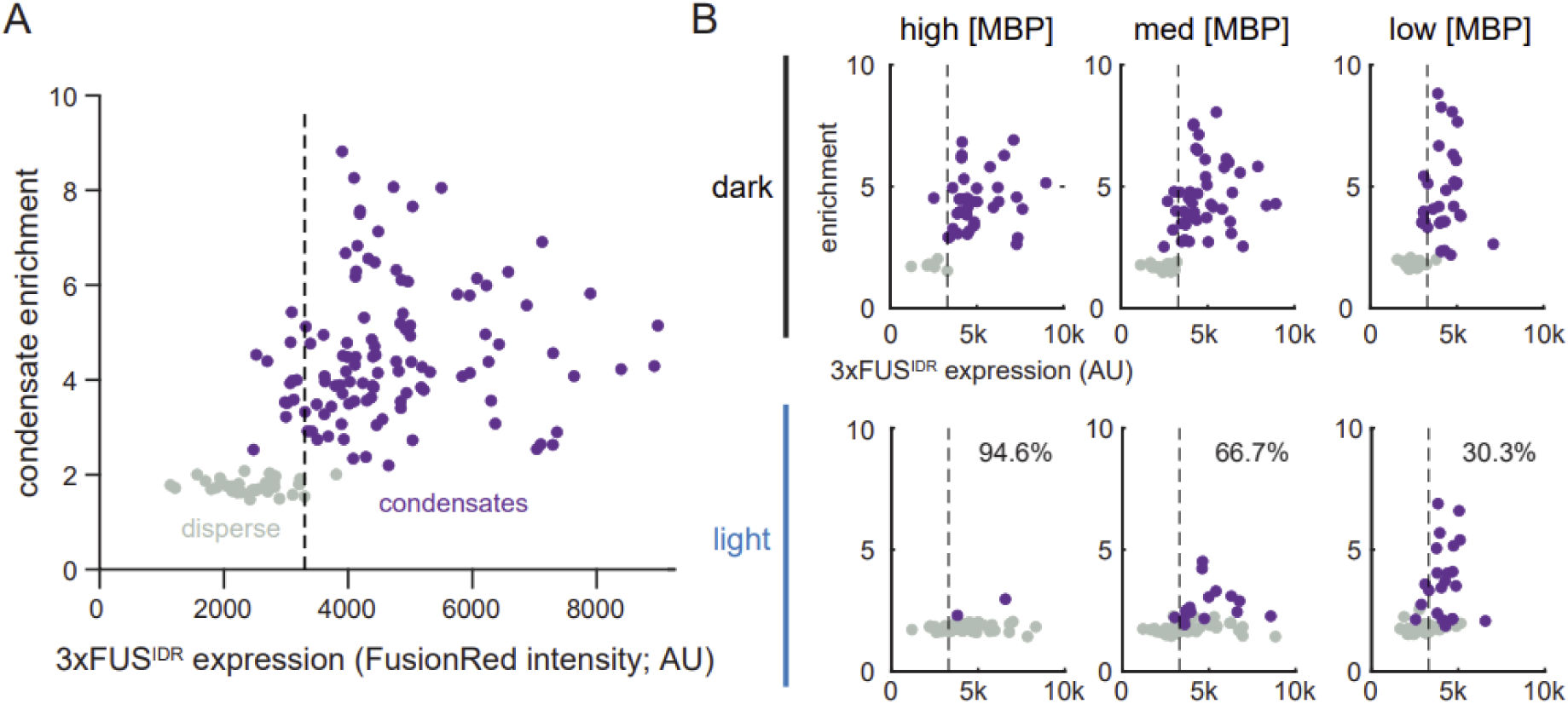
Dissolution efficiency is dependent on the concentration of MBP. **A.** Mean pixel intensity and enrichment (max/mean) in the FusionRed (3xFUS^IDR^) channel of the nucleus of an NIH3T3 cell line expressing 3xFUS^IDR^-iLID and MBP-SspB. Points are colored for cells with visible condensate formation (purple) and no resolvable condensates (gray). An approximate critical concentration is shown (dashed line). **B.** Data was binned based on the MBP concentration. The mean pixel intensity vs enrichment is shown before (top row) and after (bottom row) 3 min blue light illumination. Cells with higher concentrations of MBP showed a greater extent of condensate dissolution. Fraction of cells where condensate dissolution was observed is indicated for each MBP level.

We next set out to measure how the 3xFUS^IDR^ dissolution depended on the concentration of our OptoMBP solubilizer construct. We separately analyzed dissolution in cells with low (<3500 fluorescent units; AU), medium (3500-4500 AU), and high (>4500 AU) MBP-SspB-iRFP expression (**Fig. 3B**). Optogenetic activation of cells expressing high levels of MBP-SspB-iRFP produced rapid dissolution of 3xFUS^IDR^ condensates in nearly all cells (**Fig. 3B**, high [MBP]), whereas cells harboring medium or low MBP expression dissolved condensates in progressively fewer cells (**Fig. 3B**, med [MBP], low [MBP]). These data confirm that MBP expression level is a crucial parameter in determining the extent of optogenetic condensate dissolution. We further tested whether light-induced dissolution behaved stoichiometrically – that is, whether cells with higher 3xFUS^IDR^ expression required proportionally higher MBP expression to drive dissolution. To do so, we analyzed single-cell expression of both 3xFUS^IDR^ and MBP as well as 3xFUS^IDR^ condensation in the dark and light (**Fig. S3B**). Unexpectedly, we did not observe a clear relationship between 3xFUS^IDR^ expression level and the concentration of MBP required for light-induced dissolution. Instead, our data revealed a threshold in MBP expression (∼4500 AU) above which nearly all 3xFUS^IDR^ condensates were dissolved under blue light, regardless of their 3xFUS^IDR^ expression level. These data suggest that light-induced MBP recruitment acts cooperatively, with a switch-like change in phenotypic response observed over a narrow range of concentrations.

Taken together, these data underscore the complex nature of condensate formation and dissolution *in vivo*, wherein multiple interacting components might define parameters such as the dilute-phase concentration of each component and the threshold concentration above which condensates are observed^38^. They also point to important performance characteristics of the OptoMBP system. Above a threshold concentration of MBP, the OptoMBP system shows efficacy at reversing 3xFUS^IDR^ phase separation at a wide range of expression levels. This sharp dependence with MBP concentration but weak dependence on 3xFUS^IDR^ concentration suggests a cooperative process where a critical threshold of MBP must be surpassed to drive a robust solubilizing response. These findings may help to constrain models of the biophysical basis for MBP-induced solubilization, a process that is still poorly understood^52,61^.

### Dissolution with MBP generalizes to several condensates of interest

MBP has been reported to solubilize a wide range of target proteins in fusion constructs^50^, so we hypothesized that OptoMBP recruitment may also dissolve condensates other than 3xFUS^IDR^. We first tested other members of the FUS-EWS-TAF15 (FET) protein family, all of which form nuclear condensates, bind and regulate RNA^62^, and contain disordered domains^63^ rich in glycine, glutamine, serine, and tyrosine (**Fig. 4A**). We first created EWS and TAF15 constructs consisting of three repeats of their disordered domain, architectures that are comparable to our 3xFUS^IDR^ construct. As with 3xFUS^IDR^, 3xEWS^IDR^ and 3xTAF15^IDR^ formed condensates when transfected into HEK293T cells, and optogenetic MBP recruitment resulted in robust dissolution in each case (**Fig. 4B**). Analysis showed that CV and enrichment decreased after illumination for all three constructs (**Fig. 4C-D**).

**Fig. 4.**
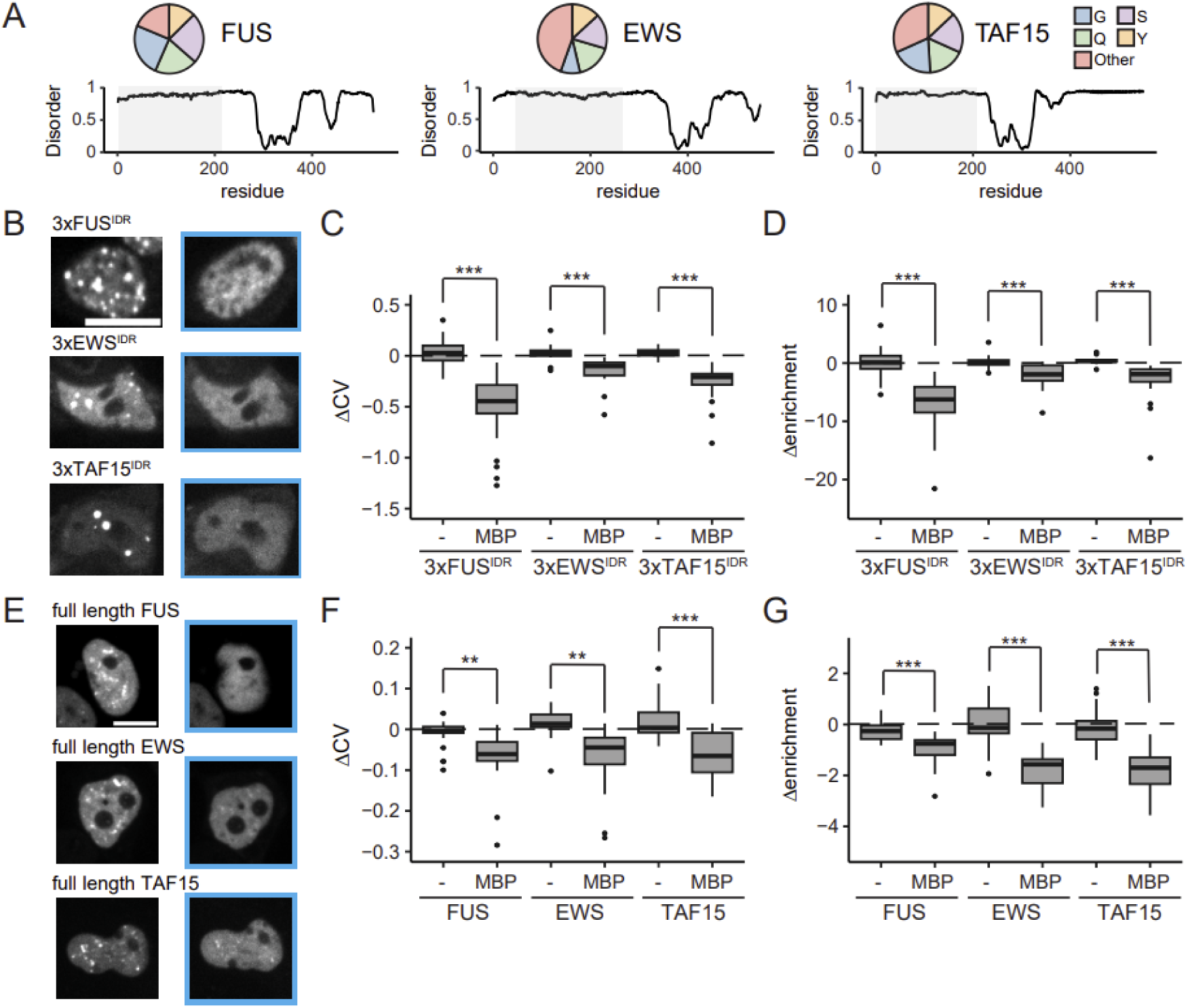
MBP-based condensate dissolution extends to FET family proteins. **A.** FUS, EWS, and TAF15 all contain an N-terminal disordered domain, gray box. This domain is highly similar amongst the three family members, consisting of over 50% glycine, glutamine, serine, and tyrosine residues. **B.** Representative images displaying nuclear condensates formed from FUS, EWS, and TAF15 constructs consisting of 3 repeats of the IDR. MBP recruitment with blue light resulted in droplet dissolution in all cases. **C.,D.** Quantification of dissolution efficiency for FET family IDRs from microscopy data by calculating the change in the coefficient of variation and enrichment after 3 min blue light stimulation. **E.** Representative images displaying nuclear condensates formed from full length FUS, EWS, and TAF15 before and after blue light stimulation to recruit MBP. **F.,G.** Quantification of dissolution efficiency for FET family full length proteins from microscopy data by calculating the change in the coefficient of variation and enrichment after 3 min blue light stimulation. Scale bars: 10 μm.

We next tested whether similar results might be obtained from the full-length FET family members containing both the disordered and the RNA-binding domains. We constructed fluorescent, iLID-tagged variants of all three FET family protein members and observed visible condensates after transient transfection, although we found that these condensates were less spherical and, for FUS and TAF15, showed lower enrichment in the condensed phase compared to their IDR-only counterparts (**Fig. S4A**). We observed dissolution in each case upon OptoMBP recruitment (**Fig. 4E**), and analysis of the changes in CV and enrichment for all FET family constructs (**Fig. 4F-G**) confirmed our qualitative observation that MBP recruitment results in dissolution.

We further probed whether MBP-based dissolution might generalize outside of the FET family to other condensate-forming proteins of interest. We tested three repeats of the disordered RGG domain from LAF-1 which has been shown to be solubilized by MBP fusion^21,22^; the full-length RNA binding protein Oskar (**Video S4**), which is required to form germ granules during *Drosophila* development^64^; mRNA decapping protein 1A (DCP1A)^38^, which participates in mRNA degradation; and PML IV, a major component of PML-nuclear bodies^65^. All four proteins formed condensates in either the nucleus (Oskar, PML IV) or cytosol (3xRGG, DCP1A) when fused to FusionRed and the iLID optogenetic recruitment domains and transfected into HEK293T cells (**Fig. S4B**). Moreover, all four proteins recruited MBP-SspB when illuminated with blue light (**Fig. S4B**, early). However, condensate dissolution was variable across this broader set of proteins. We observed robust dissolution for the 3xRGG construct and Oskar, but DCP1A and PML IV were unaffected by light-induced MBP recruitment (**Fig. S4B**, late).

Together, these results show that OptoMBP-based dissolution generalizes outside of the FET family to additional types of condensate-forming proteins in both the nucleus and cytoplasm. However, it is not universally applicable, with some condensates showing little to no dissolution upon MBP recruitment.

### Dissolution of oncogenic condensates drives transcriptional changes

Finally, we set out to apply the OptoMBP system to perturb condensate-dependent biological functions. A growing body of evidence demonstrates that condensates can represent functionally relevant states. For example, co-condensation of enzymes may enhance reaction rates^9^ or coordinate processes at length scales much longer than individual proteins^66^. Gene expression is thought to be controlled by condensation of transcription factors and other transcriptional machinery^11,32,67^, but directly testing this model by triggering condensation on demand can drive complex and unexpected outcomes. For example, while the morphogen transcription factor Bicoid has been shown to cluster at sites of active transcription^68^, light-induced clustering of the Bicoid-Cry2 optogenetic fusion protein paradoxically acts as a dominant-negative transcription factor that blocks endogenous Bicoid-induced gene expression^69^, potentially due to improper clustering between these proteins at sites other than the appropriate target enhancers. We reasoned that an alternative approach to manipulating transcription-associated condensates might be to dissolve endogenously formed condensates using the OptoMBP system, which can potentially act with high temporal and spatial resolution.

We set out to monitor the effects of OptoMBP-induced dissolution on model transcriptional condensates formed of the oncogenic fusion protein FUS-CHOP, which is associated with liposarcoma^70–72^. The FUS-CHOP oncoprotein is produced by a chromosomal translocation event wherein the disordered, N-terminal region of FUS is fused to the transcription factor CHOP (CCAAT/enhancer-binding protein homologous protein, also known as DDIT3 - DNA damage inducible transcript 3). Previous work has shown that FUS-CHOP forms biomolecular condensates and a plausible model for its action involves improper transcriptional activation triggered by FUS condensation at DNA loci targeted by the CHOP DNA binding domain. However, the role of condensate formation on the transcriptional activity of the FUS-CHOP oncoprotein is not well understood^15,73^. We developed an NIH3T3 cell line containing a doxycycline-inducible FUS-CHOP construct fused to FusionRed fluorescent protein and the iLID light activatable domain (**Fig. 5A**), which we termed FUS-CHOP/OptoMBP cells. We chose NIH3T3 cells because of their ease of manipulation and prior use in studying the transcriptional activity and oncogenic properties of FUS-CHOP and related fusion proteins^74^. We co-expressed FUS-CHOP with the rtTA transcription factor and constitutively expressed OptoMBP light-induced dissociation system (MBP-SspB-iRFP) (**Fig. 5A**). We observed nuclear FUS-CHOP-FusionRed-iLID condensates within 24 h after doxycycline induction, and, as in prior FUS-related contexts, these condensates could be rapidly and completely dissolved by light-induced MBP recruitment (**Fig. 5B-C**; **Fig. 2I-J, Video S3, Video S5**).

**Fig. 5.**
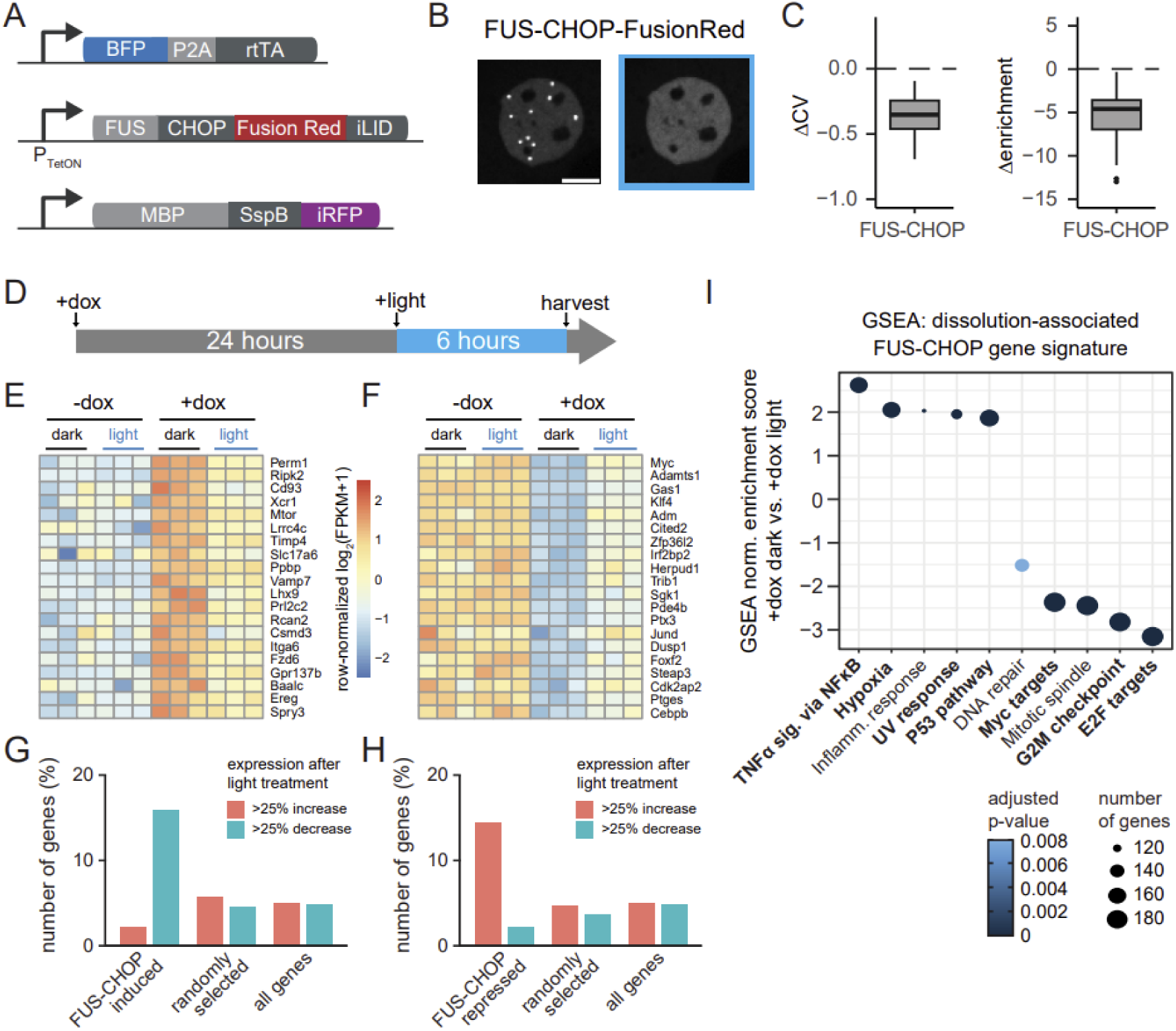
Dissolution of the oncogenic fusion protein FUS-CHOP results in reversion transcriptional changes. **A.** Schematic of approach. We generated a NIH3T3 cell line containing FUS-CHOP fused to iLID and a fluorescent marker under control of the TetOn promoter, MBP fused to SspB under control of a constitutive promoter, and rtTA under control of a constitutive promoter. **B.** Representative images showing FUS-CHOP droplet dissolution after illumination with blue light to recruit MBP. **C.** Quantification of dissolution efficiency from microscopy data by calculating the change in the coefficient of variation and enrichment of the FUS-CHOP construct after 5 min blue light stimulation. Negative values indicate condensate disruption. **D.** Experimental setup, doxycycline was added at time zero to induce expression of FUS-CHOP, after 24 hrs, blue light was applied to the cells to recruit MBP to FUS-CHOP droplets. Cells were harvested after six hours of illumination for RNASeq. **E.,F.** Heatmap displaying changes in gene expression upon dox induction of FUS-CHOP and subsequent light illumination to dissolve FUS-CHOP condensates. Rows are independently normalized and sorted based on extent of recovery to initial expression upon light stimulation. The top twenty genes among genes whose expression level increased or decreased after doxycycline induction of FUS-CHOP are shown. **G.,H.** Fraction of FUS-CHOP perturbed genes that demonstrate 25% or greater recovery after blue light illumination compared to recovery of randomly selected genes, and all genes in the analysis. **I.** GSEA analysis examining pathways that are up and down regulated in dox-induced samples after light illumination. Top five pathways up and down-regulated are shown. Scale bar: 10 μm.

We next set out to test how optogenetic FUS-CHOP dissolution might alter transcription driven by this oncoprotein. We plated FUS-CHOP/OptoMBP cells and treated them with doxycycline to induce FUS-CHOP expression for 24 h, with mock-treated cells used as a control (**Fig. 5D**). We then continued incubating cells in the dark or under 450 nm blue LED light (at an intensity of ∼1 mW/cm^2^, comparable to previously published transcriptional assays^67,75^) for an additional 6 hours to provide sufficient time to dissolve FUS-CHOP condensates and resolve any resulting changes in gene expression. Finally, we conducted bulk RNASeq to assess transcriptional changes in three replicates of each of the four conditions (**Fig. 5E-F**).

We first assessed the transcriptional consequences of FUS-CHOP expression by identifying differentially expressed genes between doxycycline-treated and untreated conditions in the dark (**Fig. S5A**). Differential expression analysis revealed widespread expression changes with 283 genes increasing more than 2-fold after doxycycline induction and 570 genes decreasing more than 2-fold. We compared these data to a previously published FUS-CHOP RNASeq data set in a human fibrosarcoma cell line, HT1080^76^; looking at the top 100 hits in HT1080 published genes that also displayed at least a 2-fold change in expression in our study, ∼80% displayed the same direction of change in both datasets (**Fig. S5B**).

We next assessed gene expression differences between dark and blue light in the absence of doxycycline to control for any illumination-induced changes in gene expression that were not related to FUS-CHOP or our optogenetic system (**Fig. S5C**). Light-triggered gene expression that is independent of optogenetic tool expression has been previously documented in neuronal contexts, albeit at the higher illumination intensities (>100 mW/cm^2^) than our system requires^77^.

To minimize confounding effects of illumination on our analysis, we implemented stringent p-value (<0.05) and fold-change (>25%) cutoffs to exclude light-controlled genes from subsequent analysis. This analysis yielded 253 genes whose expression differed upon blue light illumination in the absence of FUS-CHOP. We also performed gene set enrichment analysis (GSEA) comparing non-illuminated and illuminated samples in the absence of doxycycline. This analysis revealed enrichment for genes involved in the unfolded protein response, particularly heat shock response factors (**Fig. S5D**).

Finally, we determined how the signature of FUS-CHOP gene expression was altered in response to light-induced dissolution by the OptoMBP system. We hypothesized that genes whose expression is altered by FUS-CHOP induction might revert towards their original expression level in response to dissolution by OptoMBP. To test this hypothesis, we compared the changes in expression of FUS-CHOP-responsive genes to randomly chosen gene sets drawn from the same distribution of expression levels in our dataset, as well as to the overall changes across all genes (**Fig. S5E-G**). We found that light stimulation disproportionately reverted gene expression from a FUS-CHOP perturbed state back toward basal expression. Approximately 15% of FUS-CHOP-upregulated genes were reduced by at least 25% by light, compared to 5% of randomly chosen genes; only 2% of FUS-CHOP-upregulated genes showed a further increase upon illumination (**Fig. 5G**). Similar proportions were also observed for genes whose expression was decreased upon FUS-CHOP expression and restored to higher levels upon illumination (**Fig. 5H**). These data indicate that even a short 6 h light stimulus is sufficient to partially revert FUS-CHOP-induced changes in gene expression.

To further characterize the genes affected by light-induced dissolution, we performed gene set enrichment analysis (GSEA) on FUS-CHOP-expressing cells before and after illumination (**Fig. 5I**). We found that this gene set was enriched for various gene categories: NFκB, pro-inflammatory, and p53 gene signatures, as well as de-enrichment for Myc targets, E2F targets, and G2/M associated gene expression. These gene sets also closely matched a prior study examining the consequences of FUS-CHOP silencing in a liposarcoma cell line^78^ (**Fig. 5I**; bold labels), suggesting that dissolution of transcriptional condensates functionally resembles overall loss of the transcription factor. Taken together, our results suggest that FUS-CHOP broadly alters gene expression within 24 h, and that these effects can be partially reversed by specifically targeting FUS-CHOP phase separation with OptoMBP. These data are consistent with a direct role for FUS-CHOP condensation in promoting gene expression, as disassembly of FUS-CHOP condensates began to revert gene expression changes even under continued induction of oncoprotein expression.

## Discussion

In this study, we report the first optogenetic system to dissolve constitutively formed biomolecular condensates. Optogenetic tools offer many advantages over other strategies, including the potential for rapid reversibility, fine spatial control, and a high degree of specificity^79^. We found that light-based recruitment of the classic solubilizer, maltose-binding protein (MBP) was highly effective at dissolving a variety of protein condensates: those formed from FET family members, the Oskar RNA binding protein, the LAF-1 RGG domain, and the FUS-CHOP oncoprotein. Light-based MBP recruitment can be quite potent, even matching the efficacy of the broad-spectrum condensate disrupter 1,6-hexanediol. Moreover, optogenetics confers additional advantages in reversibility and spatial control. For example, one can envision future studies where MBP is used to dissolve transcription factor condensates at a single transcriptionally active locus of interest to specifically alter their presence or composition and monitor real-time effects on gene expression or genome spatial organization.

Our results also raise an open question: what is the mechanism by which MBP achieves its solubilizing effects? One can imagine several possibilities: MBP might be of sufficient size to sterically block favorable interactions within the condensate; MBP could preferentially bind the condensate-forming protein in the dilute phase, thereby destabilizing condensates through polyphasic linkage^80^; the high solubility of MBP may alter the saturation concentration of the complex; or MBP may bind to the same sequences that would normally be used to form multivalent interactions. In general, we do not observe a correlation between cargo size and dissolution propensity, suggesting that the first of these mechanisms (steric hindrance) is unlikely to be the cause of MBP solubilization, although the other proposed mechanisms cannot be excluded. Surprisingly, we observe a relatively sharp threshold in MBP concentration at which condensate dissolution is observed, and this threshold does not appear to vary in proportion to the concentration of the target protein, suggesting that a simple stoichiometric model of MBP-target binding is not sufficient to explain the effect (**Fig. 3**). Despite extensive study, the mechanism by which MBP solubilizes target proteins in direct fusion constructs is not fully understood^50,52,61^. One could imagine OptoMBP serving as a platform to study the mechanism of MBP solubilization. There is still much to be done to decipher the code that links condensate and solubilizer.

Our study establishes that the OptoMBP system is exceptionally effective at dissolving condensates formed of FET family proteins. This includes isolated FET intrinsically disordered domains, full-length family members that retain RNA binding domains, and an oncogenic fusion protein containing the FUS IDR and a DNA binding domain. In the latter case, we were further able to examine the link between phase separation of a transcription regulator (FUS-CHOP) and downstream gene expression. We found that OptoMBP-based dissolution partially reversed FUS-CHOP-driven gene expression changes within 6 h of illumination, mimicking the gene signature obtained by directly silencing FUS-CHOP^78^. While prior studies established that fusion to the FUS domain controls both phase separation and oncogenicity^73,74^, our results hint at a causal link between FUS-CHOP material state and its function as a transcription factor. We anticipate that similar approaches could be useful to better understand the role of transcription factor condensates in other gene expression contexts where condensation may be as likely to interfere with gene expression as it is to promote it^69,81^.

The OptoMBP system is an important initial step towards programmable, optogenetic dissolution of any condensate. In future work, it will be essential to better define the mechanism of MBP-induced solubilization to be able to predict which condensates will be dissolved by OptoMBP and to rationally design improved variants that expand its reach to additional protein aggregates. It will also be important to develop systems that enable MBP recruitment to endogenous protein targets, not just those which are tagged with a dimerization domain. The recent development of light-controlled nanobody and monobody binders could solve this problem, enabling light-gated binding to endogenous targets^82,83^. Nevertheless, the OptoMBP system adds new functionality to the toolbox of approaches to manipulate biomolecular condensates and regulate their biological functions.

## Materials and Methods

### Plasmid Construction

Constructs utilized in this study were inserted in a pHR or pPB vector that included a SFFV or TetOn promoter and WPRE terminator. Constructs were cloned with InFusion (Clontech) using backbone PCR. The MBP and GST genes were a gift from Prof. Daniel Hammer, University of Pennsylvania. RGG, DCP1A, and PMLIV genes were a gift from David Sanders and Prof. Cliff Brangwynne, Princeton University. The Oskar gene was a gift from Prof. Elizabeth Gavis, Princeton University. EWS, TAF15, and CHOP genes were generated by gene synthesis (Integrated DNA Technologies).

### Cell transient transfection

For microscopy experiments, HEK293T cells were plated on a glass-bottom 96-well plate (Cellvis) at around 30% confluency. The next day, these cells were transfected using lipofectamine 3000 (ThermoFisher) following the manufacturer’s instructions. For experiments involving two constructs, equal amounts of both plasmids were co-transfected. Cells were imaged 24 hours after transfection.

### Lentivirus production and transduction

Lenti-X cells were plated at approximately 30% confluency in a 6 well plate. A pHR expression plasmid and lentiviral packaging plasmids were co-transfected using the Fugene HD transfection reagent (Promega). 52 hours after transfection, the supernatant was collected and passed through a 0.45 μm filter. Cells to be infected with lentivirus were plated at 30% confluence in a 6 well plate and 500μL of virus solution, 50μL of 1M HEPES, and 1μL of 5 μg/mL polybrene (Sigma Aldrich) were added. Cells were then expanded for sorting.

### Stable cell line generation

To generate stable NIH3T3 cell lines co-expressing 3xFUS^IDR^-FusionRed-iLID and MBP-SspB-iRFP, we produced lentivirus vectors for both genes. NIH3T3 cells infected with MBP-SspB-iRFP lentivirus and iRFP positive cells were isolated using Fluorescence-activated Cell Sorting (FACS). These MBP-expressing cells were then infected with 3xFUS^IDR^-FusionRed-iLID lentivirus. iRFP and FusionRed double positive cells were then isolated with FACS. To generate FUS-CHOP cell lines, we first generated a cell line expressing rtTA. NIH3T3 cells were infected with BFP-P2A-rtTA lentivirus and BFP positive cells were isolated. To these cells, a piggybac transposon plasmids containing both MBP-SspB-iRFP and TetOn-FUS-CHOP-FusionRed-iLID genes were cotransfected with the piggybac transposease. Cells were then sorted for expression of both BFP and iRFP. Sorting was performed on a SH800S Cell Sorter (Sony Biotechnology).

### Cell imaging

Confocal microscopy was performed on a Nikon Eclipse Ti microscope equipped with a Yokogawa CSU-X1 spinning disk, Agilent 4 color laser line module, iXon DU897 EMCCD camera, and Mightex Polygon400 digital micromirror device (DMD). Live cells were maintained at 5% CO_2_ and 37°C during imaging and imaged using a 60X oil immersion objective. For dissolution experiments images were taken with the 561 nm and 650 nm laser lines.

### Chemical and small molecule treatment

To dimerize FRB and FKBP, rapalog AP21967 (Takara) was added to a final concentration of 500 nM. To induce expression of FUS-CHOP, doxycycline was added to a final concentration of 1 μg/mL. To fully dissolve droplets, 1,6-hexanediol was dissolved in DMEM and added to a final concentration of 10%.

### Blue light activation

To dimerize iLID and SspB, 450 nm light was applied with an LED and delivered to the region being imaged through a DMD between acquisitions. For subcellular activation, the DMD region of interest was adjusted to encompass about half of the cell being imaged or encompass a single condensate in the cell nucleus. DMD was set to 50% power, 50% dithering for illumination experiments, except subcellular illumination where the DMD was reduced to 10% power to reduce off-target light stimulation.

### Data quantification and analysis

Microscopy images were quantified using ImageJ (NIH) and MATLAB R2022a (MathWorks). In ImageJ, the mean, max, and standard deviation of the red channel pixel intensity in the nucleus (FUS, oskar, PML IV) or cytoplasm (RGG, DCP1A) was measured. ΔCV was defined as (𝜎_f_/μ_f_)-(𝜎_i_/μ_i_) and Δenrichment was defined as (max_f_/μ_f_)-(max_i_/μ_i_). Plots were created using RStudio (Posit) or MATLAB. Significance was determined using Student’s t-test: n.s.; p > 0.05 n.s., *;p < 0.05, **;p<0.01, ***;p<0.001.

### RNASeq

Total RNA was isolated from our FUS-CHOP NIH3T3 cell line using the RNeasy mini kit (Qiagen). RNA-Seq directional library preparation and sequencing on Illumina NovaSeq 6000 was performed by the Princeton Genomics Core Facility. Using Galaxy^84^, reads were mapped using the Spliced Transcripts Alignment to a Reference (STAR) algorithm^85^ and then counted using the featureCounts algorithm^86^. Differential expression analysis was performed using DESeq2^87^ in RStudio. Volcano plots were generated with the EnhancedVolcano R package using and heatmaps were generated using the pheatmap R package. Gene set enrichment analysis (GSEA) was performed in RStudio with FGSEA^88^ using the mouse-ortholog hallmark gene sets^89^.

## Acknowledgements

We thank all members of the Toettcher and Aoki laboratories for helpful discussions and expertise. We thank the Princeton Genomics Core for RNA sample library preparation/sequencing and guidance. This work was supported by the International Research Collaboration Center at the National Institute of Basic Biology, Japan (to E.H.B.R.) as well as NIH grant U01DK127429 and a Vallee Scholar award (to J.E.T.).

## Supplementary Information

**Fig. S1.**
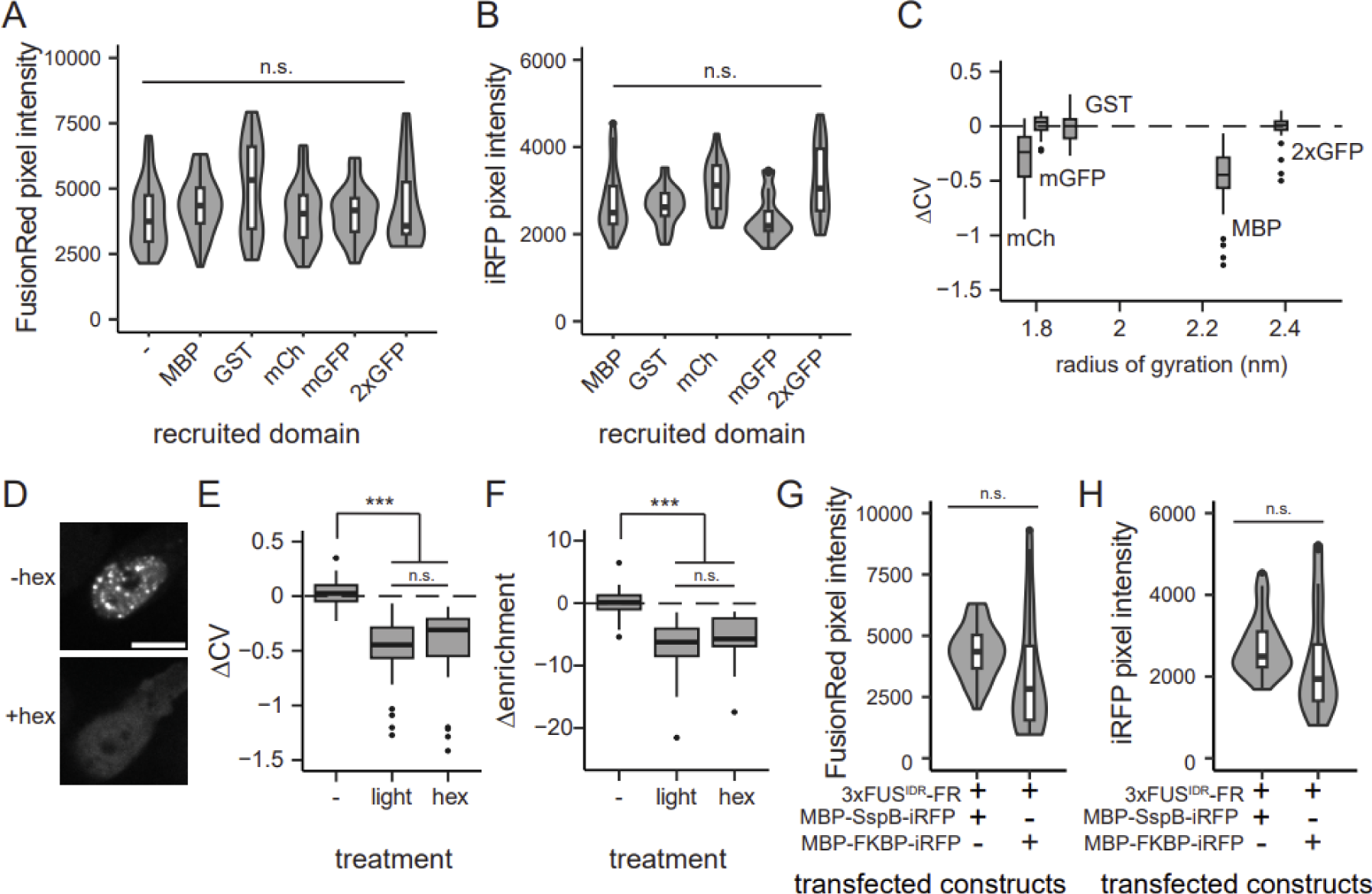
**A.,B.** Comparison of mean pixel intensity in the FusionRed (3xFUS^IDR^) and iRFP (solubilizer candidate) channels between cells used to compare dissolution efficacy of candidate solubilizer proteins. **C.** Comparison of the size of recruited protein and dissolution efficiency shows no clear trend. **D.** Representative images displaying dissolution of 3xFUS^IDR^ condensates using 1,6-hexanediol. **E.,F.** Change in CV and enrichment of 3xFUS^IDR^ 3 min after addition of 1,6-hexanediol. MBP recruitment induced droplet dissolution was equally as effective as 1,6-hexanediol induced droplet dissolution. **G.,H.** Comparison of mean pixel intensity in the FusionRed (3xFUS^IDR^) and iRFP (MBP) channels between cells used to compare dissolution efficacy between optogenetic and chemogenetic recruitment. Scale bar: 10 μm.

**Fig. S2.**
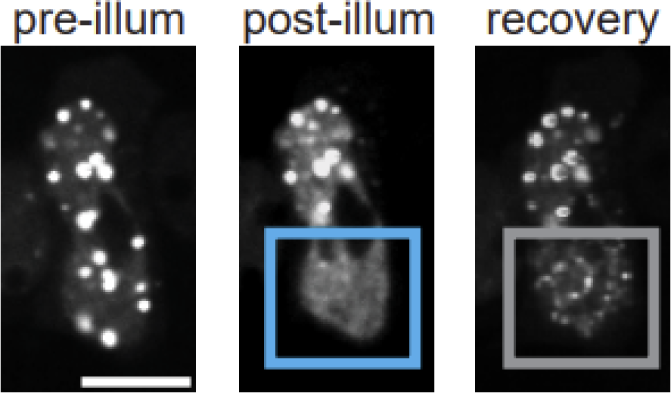
Images displaying 3xFUS^IDR^ droplet dissolution in an illuminated region of a cell after 3 min light stimulation and recovery in the dark state for 7 mins. A disparity in condensate size between illuminated and non-illuminated regions persists after the removal of blue light.

**Fig. S3.**
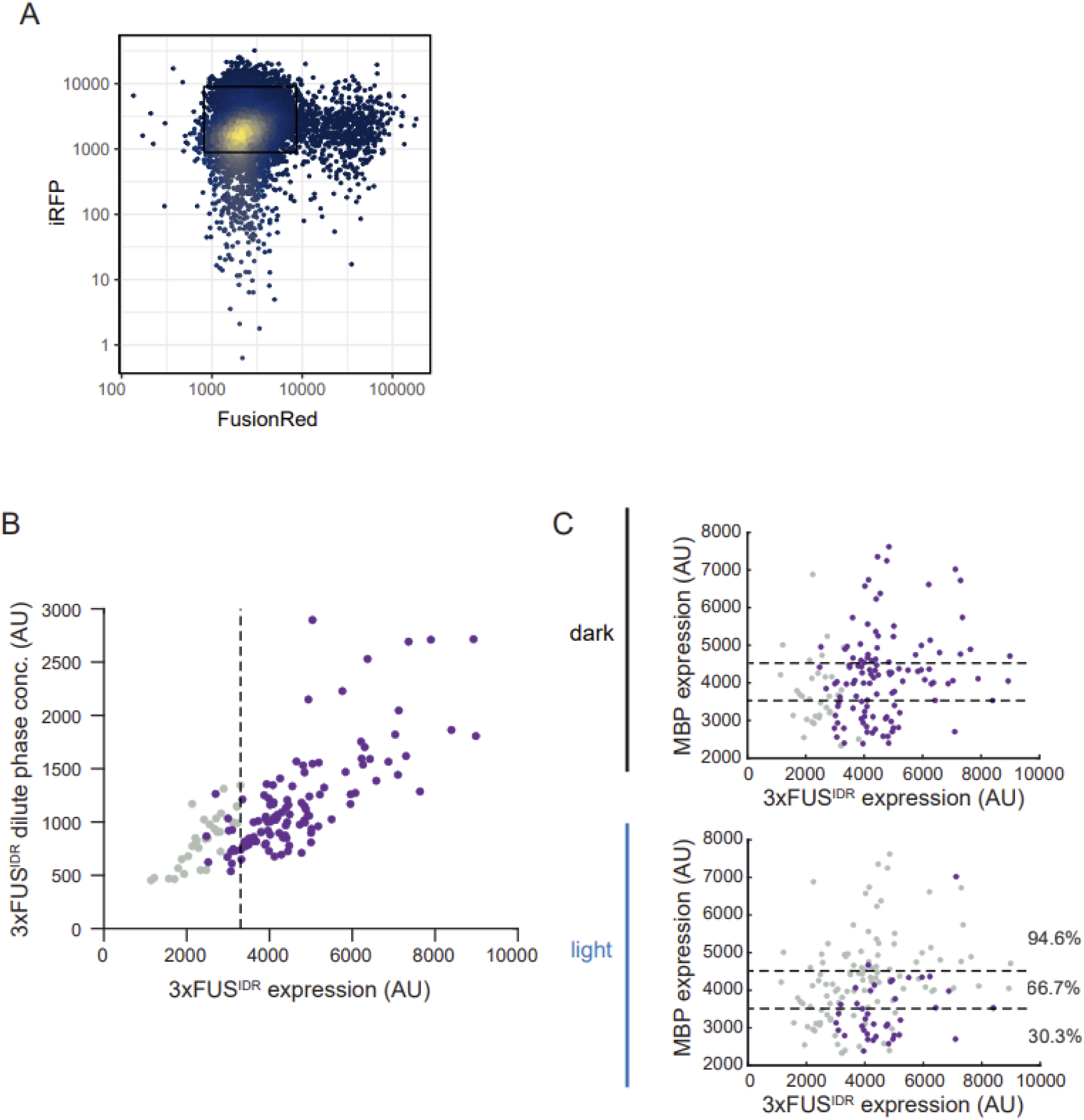
**A.** Flow cytometry data showing FusionRed (FUS^IDR^) and iRFP (MBP) expression. Cells in the boxed area were sorted for microscopy experiments. **B.** Comparison between the mean FusionRed pixel intensity (proxy for overall FUS^IDR^ concentration) and the minimum pixel intensity (proxy for FUS^IDR^ concentration in the dilute phase). The approximate critical concentration is shown (dashed line). Above the critical concentration, the dilute phase concentration of FUS^IDR^ continues to rise, indicating that 3xFUS^IDR^ condensates are likely multiphase condensates. **C.** Comparison of MBP and FUS^IDR^ expression before (top) and after (bottom) 3 min blue light illumination. Cells with visible droplets are displayed as purple.

**Fig. S4.**
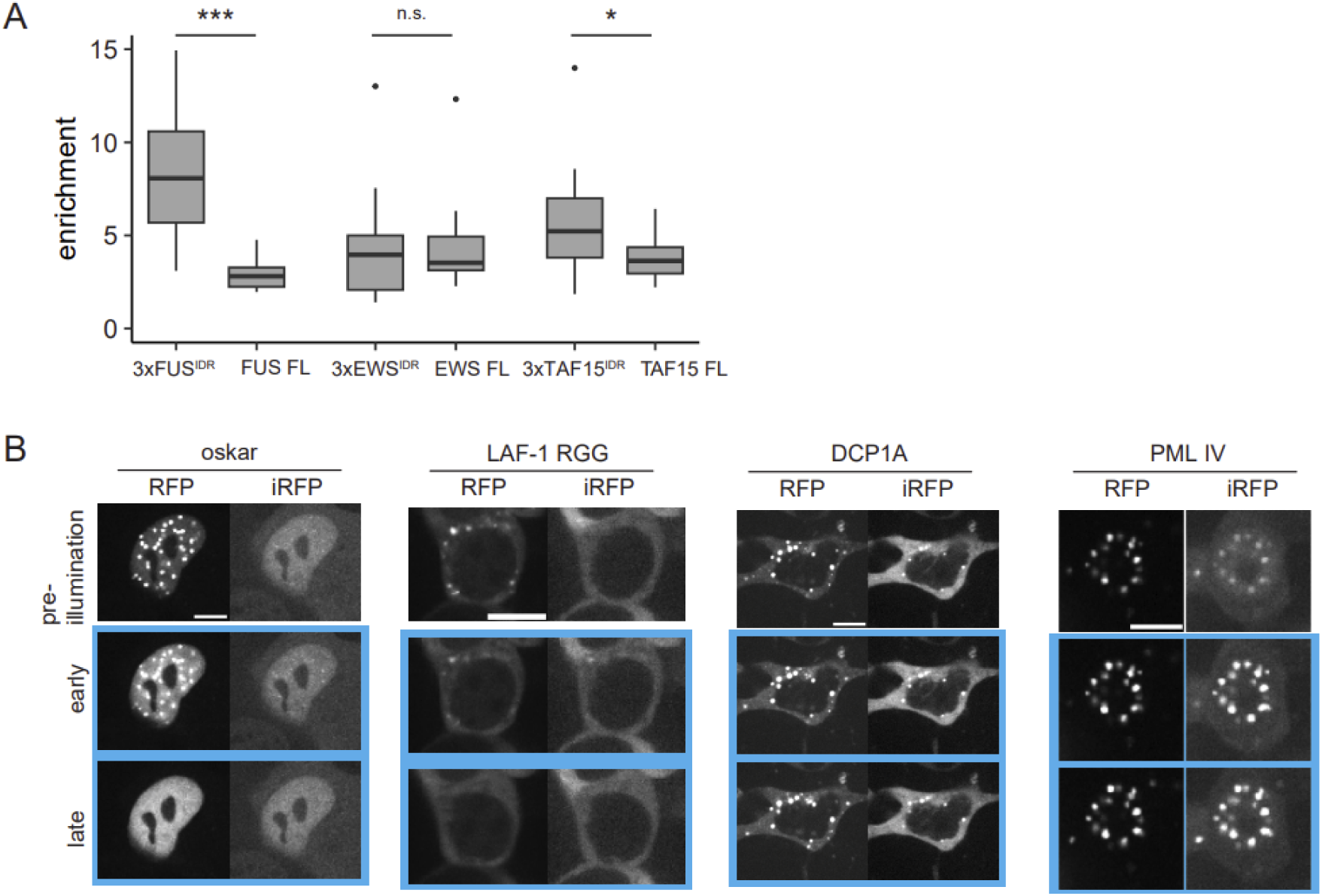
**A.** Comparison of dark state droplet enrichment for FET family IDRs and full length proteins. **B.** Representative microscopy images of cells expressing phase separating proteins (oskar, LAF-1 RGG, DCP1A, PML isoform IV) in the dark state, shortly after blue light illumination (early), and 3 minutes post illumination (late). FusionRed (condensate forming protein) and iRFP (MBP) channels are shown.

**Fig. S5.**
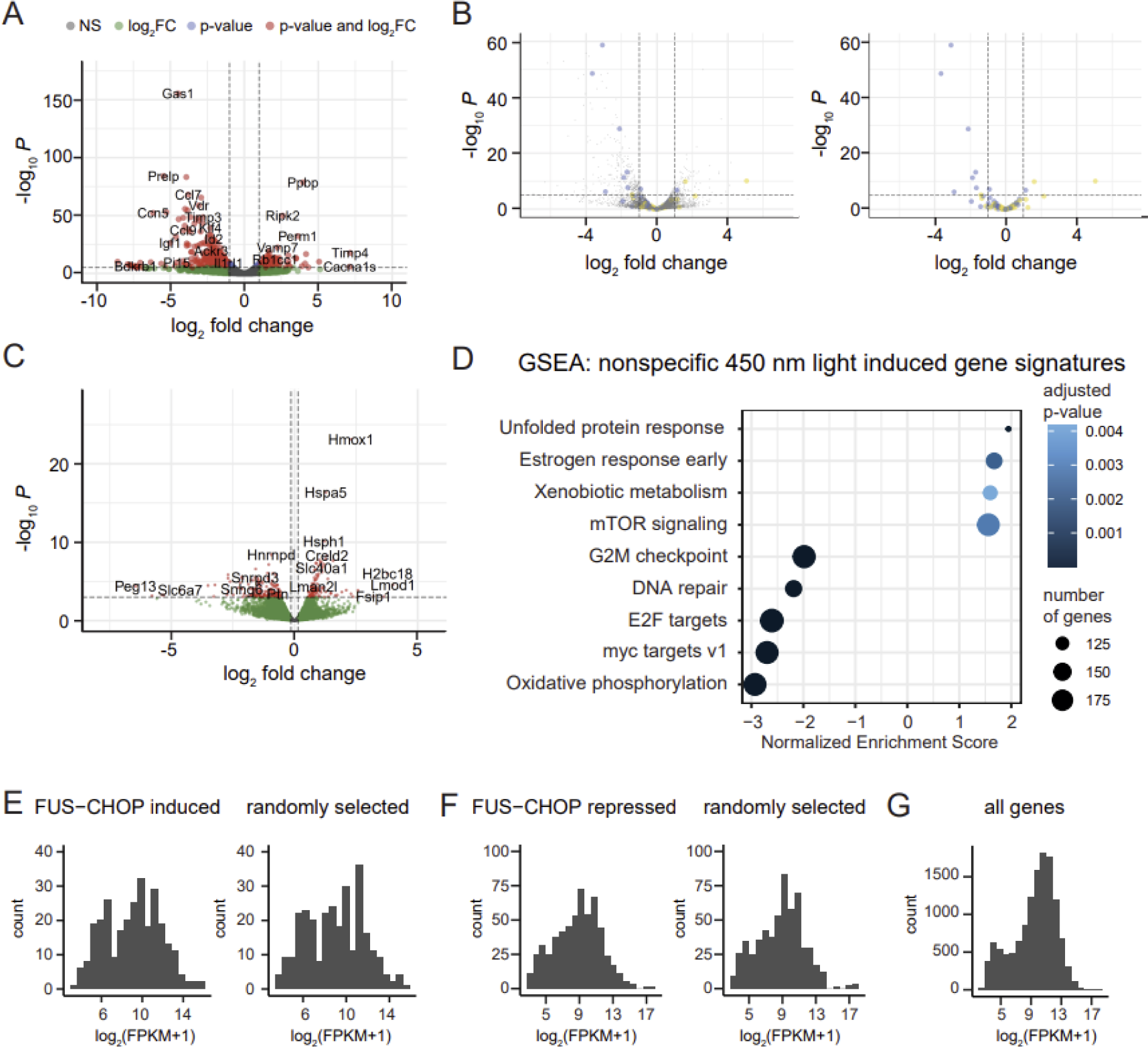
**A.** Volcano plot displaying changes in gene expression after addition of doxycycline to induce expression of FUS-CHOP. **B.** Top hits from previously published FUS-CHOP RNASeq dataset. Upregulated genes are shown in yellow, downregulated are shown in blue. **C.** Volcano plot displaying changes in gene expression after light stimulation, in the absence of doxycycline to identify blue light perturbed genes. **D.** GSEA analysis showing pathways that are affected by blue light stimulation.**E.,F.,G.** Histograms showing normalized gene counts for FUS-CHOP induced, repressed, and randomly selected genes.

**Video S1.** Time-lapse imaging of a HEK293T cell nucleus expressing

3xFUS^IDR^-FusionRed-FRB-iLID and MBP-SspB-iRFP. FusionRed channel is shown with the blue box indicating times when 450 nm light is applied to the sample to dimerize iLID/SspB and recruit MBP, resulting in dissolution. Related to Figure 2D-F.

**Video S2.** Time-lapse imaging of a HEK293T cell nucleus expressing

3xFUS^IDR^-FusionRed-FRB-iLID and MBP-SspB-iRFP. FusionRed channel is shown with the blue box indicating times when 450 nm light is applied to the bottom half of the cell only to dimerize iLID/SspB and recruit MBP, resulting in dissolution. Related to Figure 2G-H.

**Video S3.** Time-lapse imaging of a NIH3T3 cell nucleus expressing

FUS-CHOP-FusionRed-iLID and MBP-SspB-iRFP. FusionRed channel is shown with the blue box indicating the region where blue light was applied to induce dissolution. Related to Figure 2I-J.

**Video S4.** Time-lapse imaging of a HEK293T cell nucleus expressing oskar-FusionRed-iLID and MBP-SspB-iRFP. FusionRed channel is shown with the blue box indicating times when 450 nm light is applied to the sample to dimerize iLID/SspB and recruit MBP. Related to Figure S4B.

**Video S5.** Time-lapse imaging of a NIH3T3 cell nucleus expressing

FUS-CHOP-FusionRed-iLID and MBP-SspB-iRFP. FusionRed channel is shown. Blue light is applied throughout to dimerize iLID/SspB and recruit MBP to the condensates. Related to Figure 5B.

